# High-throughput live-imaging of embryos in microwell arrays using a modular, inexpensive specimen mounting system

**DOI:** 10.1101/111583

**Authors:** Seth Donoughe, Chiyoung Kim, Cassandra G. Extavour

## Abstract

Live-imaging embryos in a high-throughput manner is essential for shedding light on a wide range of questions in developmental biology, but it is difficult and costly to mount and image embryos in consistent conditions. Here, we present OMMAwell, a simple, reusable device that makes it easy to mount up to hundreds of embryos in arrays of agarose microwells with customizable dimensions and spacing. OMMAwell can be configured to mount specimens for upright or inverted microscopes, and includes a reservoir to hold live-imaging medium to maintain constant moisture and osmolarity of specimens during time-lapse imaging. All device components can be cut from a sheet of acrylic using a laser cutter. Even a novice user will be able to cut the pieces and assemble the device in less than an hour. At the time of writing, the total materials cost is less than five US dollars. We include all device design files in a commonly used format, as well as complete instructions for its fabrication and use. We demonstrate a detailed workflow for designing a custom mold and employing it to simultaneously live-image dozens of embryos at a time for more than five days, using embryos of the cricket *Gryllus bimaculatus* as an example. Further, we include descriptions, schematics, and design files for molds that can be used with 14 additional vertebrate and invertebrate species, including most major traditional laboratory models and a number of emerging model systems. Molds have been user-tested for embryos including zebrafish (*Danio rerio*), fruit fly (*Drosophila melanogaster*), coquí frog (*Eleutherodactylus coqui*), annelid worm (*Capitella teleta*), amphipod crustacean (*Parhyale hawaiensis*), red flour beetle (*Tribolium castaneum*), and three-banded panther worm (*Hofstenia miamia*), as well mouse organoids (*Mus musculus*). Finally, we provide instructions for researchers to customize OMMAwell inserts for embryos or tissues not described herein.

**Summary Statement:** This *Techniques and Resources* article describes an inexpensive, customizable device for mounting and live-imaging a wide range of tissues and species; complete design files and instructions for assembly are included.

## Introduction

Live-imaging embryos and small organisms in a repeatable, high-throughput manner is crucial for understanding the cellular dynamics that underlie the development of multicellular bodies (Farhadifar et al., 2015; Kuntz & Eisen, 2014). High-throughput imaging allows one to assess subtle phenotypes that can arise from functional genetics experiments, study standing variation within a population, and understand the role of noise in developmental processes. To that end, some research groups have turned to microfluidic devices (Chronis, 2010; Cornaglia et al., 2015; Crane, Chung, Stirman, & Lu, 2010; Wielhouwer et al., 2011). Such microfluidic apparatuses can be constructed to perform precise and complex experimental manipulations, but designing and fabricating these devices is a laborious process. For the purpose of imaging embryos, another option is to fabricate a custom mold that can be used to cast an agar or agarose microwell array. Molds can be milled from plastic (Kainz, 2009) or aluminum (Herrgen, Schroter, Bajard, & Oates, 2009), or 3D-printed (Alessandri et al., 2017; Gregory & Veeman, 2013; Wittbrodt, Liebel, & Gehrig, 2014). Although these techniques are effective, in each case it is expensive and/or time-consuming to prototype a single mold design.

To address this outstanding need, we developed OMMAwell (**O**pen **M**odular **M**old for **A**garose Micro**well**s), an all-in-one device that allows the user to swap out any number of customized mold inserts. These inserts can be prototyped quickly and cheaply, requiring only a sheet of acrylic plastic and a laser engraver. These mold inserts lock into the device, which can be configured in several ways to optimally mount specimens for upright and inverted microscopes. Using this tool, we can mount up to hundreds of embryos at once in a microwell agarose array, keing track of each embryo by its position in the array, and then efficiently image them with uniform illumination. The modular mold inserts can be exchanged to alter the size, shape, orientation, and spacing of microwells. OMMAwell is therefore adaptable for different experimental designs or even diverse species.

As an example case, we demonstrate a workflow for making a custom mold insert for embryos of the cricket *Gryllus bimaculatus*.These cricket embryos can be imaged through their transparent eggshells. During previous efforts to live-image embryonic development within the eggs – using confocal and widefield microscopy – only a few embryos could be imaged at a time, and the mounting process was inconsistent and time-intensive (Donoughe & Extavour, 2016; Nakamura et al., 2010). Eggs were either manually glued to a coverslip one at a time (Nakamura et al., 2010) or placed in blocks of rubber polymer in which troughs had been hand-cut with a razor (Donoughe & Extavour, 2016). Mounting is similarly laborious for most animal laboratory models, which limits the sample size of experiments and reduces reproducibility. However, we show that with OMMAwell, it is straightforward to mount of dozens or hundreds of embryos in a manner that is suitable for 2D or 3D long-term time-lapse recordings.

In the Supplemental Materials, we include detailed instructions for assembling the OMMAwell mounting device, plus suggestions for modifying the device to suit the particular requirements of any desired model system. We have also designed and beta-tested mold inserts for embryos of 13 additional species, including zebrafish, fruit fly, frog, annelid worm, amphipod crustacean, red flour beetle, and three-banded panther worm, as well as mouse neurospheres. Descriptions, schematics, and design files for all of these mold inserts are provided.

## Results and Discussion

To design the first iteration of a cricket embryo mold, we collected and measured dimensions of freshly laid eggs (Figure 1A). The eggs are roughly ellipsoidal in shape, 2500 to 3200 µm in length, and 475 to 650 µm in width (Figure 1B, C). We designed the mold insert to have rectangular posts 2930 µm long by 570 µm wide, each of which will create an agarose microwell able to snugly accommodate the majority of eggs (Figure 1D).

**Figure 1.**
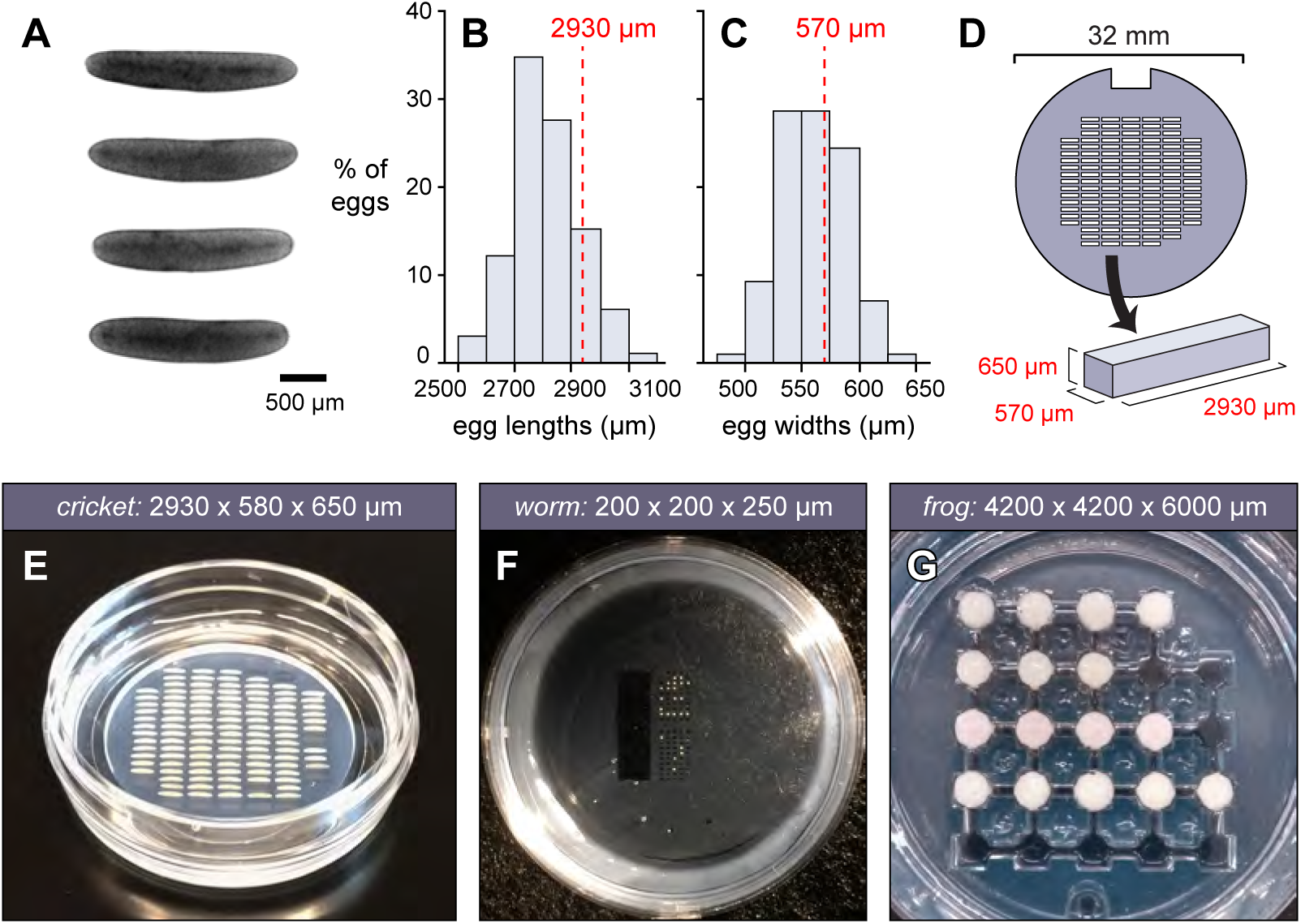
Designing microwells to hold cricket eggs. Freshly laid cricket eggs (A) were measured and their lengths (B) and widths (C) plotted (n = 98; scale bar = 500 μm). Based on the size distribution, we chose dimensions of 2930 x 570 x 650 μm (embryo length x width x height), which were values such that approximately 75% of eggs would fit into the troughs. In practice, because the wells are made of agarose, more than 95% of eggs fit into these wells. (D) Raised posts of those dimensions were formed by engraving an acrylic insert. 120 such posts were arranged in a grid. (E) Agarose microwells made using this insert, loaded with cricket eggs. (F, G) Example molds for an annelid worm and coqui frog, with microwell dimensions listed. Photos in F and G by Elaine Seaver (The Whitney Laboratory for Marine Bioscience) and Mara Laslo (Harvard University) respectively.

The “mold insert” is the only piece of the OMMAwell that must be tailored to create wells of appropriate dimensions for one’s samples of interest. To make the cricket mold insert, the inverse of our desired pattern was laser-engraved into acrylic to a depth of 650 μm. This is deep enough to contain the embryo, but close enough to the surface to be imaged within the working distance of 5x and 10x microscope objectives. The microwells were arranged into a truncated grid pattern that fit within a 26 mm circle, so that all microwells couldbe viewed through the circular 27 mm in diameter coverslip (surface area 531 mm^2^) of a 35 mm glass-bottom petri dish (Figure 1D, E; see Supplemental Materials S1). Given the dimensions of these particular embryos, we were able to fit 120 wells into the grid. For embryos of different dimensions, more or fewer wells may be able to fit into the coverslip field (e.g. 24 wells for the coquí frog *Eleutherodactylus coqui*; 294 wells for the fruit fly *Drosophila melanogaster;* see Supplemental Materials S1). One post was omitted in the corner, to make it possible to unambiguously orient the dish.

The resulting microwells for cricket embryos are shown in Figure 1E, alongside example microwells for two additional species, the three-banded panther worm (Figure 1F) and the coquí frog (Figure 1G). Note that in the latter two cases, the mold microwells were not simple rectangles, but instead had more complex shapes. Since the mold inserts are designed in 2D using a simple drawing program (see Supplemental Materials S2), it is easy for a user without prior experience to design a complex custom mold. Details for all 14 user-tested mold inserts are in Supplemental Materials S1. When testing a new mold insert, it was typically necessary iterate the design several times. An advantage of using a laser engraver to make the mold inserts is that design and fabrication is extremely rapid. In our hands, pending embryo availability, it was possible to design, fabricate, and test up to five successive iterations of a mold insert in one day.

With the mold insert ready, we cut and assembled the non-customized OMMAwell components. Detailed assembly instructions with photo guides are in Supplemental Materials S3; the design file for each component is included in Drawing Exchange Format (DXF) in Supplemental Materials S2. These files can be opened and edited by many design or drawing software packages, including AutoCAD, FreeCAD, Solidworks, SketchUp Pro, Adobe Illustrator, and CorelDRAW. All of the pieces can be made from a single sheet of 6 mm thick acrylic sheet on a laser cutter. Alternatively, if the user does not have access to a laser cutter, the components can be fabricated by online providers of laser cutting services.

For a single mold insert, there are three possible OMMAwell configurations, each of which is useful for different purposes (Figures 2 and 3). Below we discuss the usage of each configuration separately.

**Figure 2.**
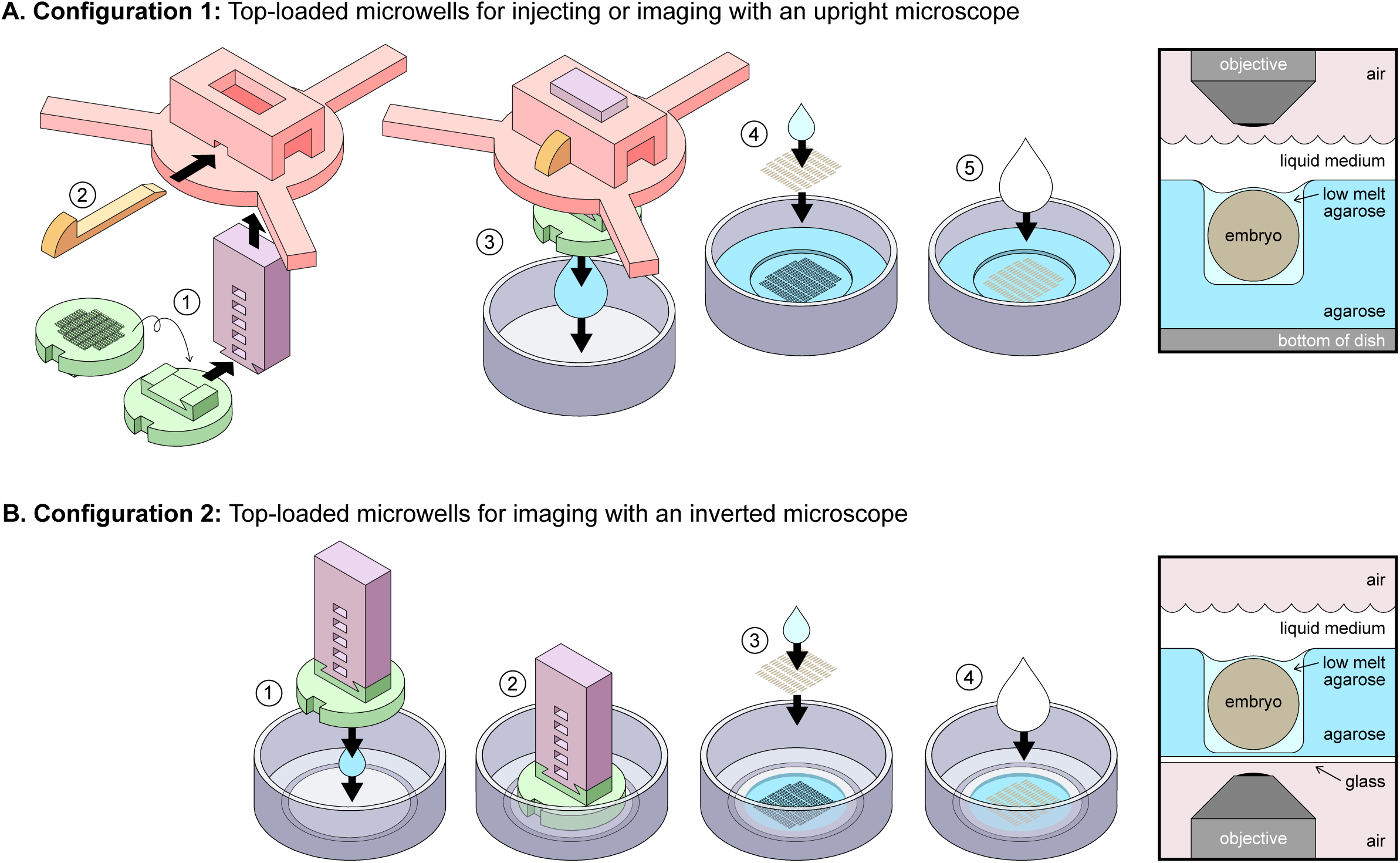
OMMAwell configurations for top-loaded microwells. (A) *Configuration 1: Top loaded microwells for injecting or imaging with an upright microscope.* 1) The mold insert (green) is inverted and connected to the slide (purple), which is placed into the upright platform (pink). 2) After the desired height is chosen, the pin (orange) is inserted. 3) Molten agarose is poured into a plastic petri dish and the mold assembly is lowered into it. 4) After the agarose sets, the mold insert is removed and eggs are placed into the wells, either individually with forceps or many at once by transferring the eggs in water with a cut plastic pipette. Excess water is removed by pipet and then wicked away with piece of lint-free lens paper. Then, 40-100 μL of molten low-melt agarose, kept at 42°C, is added to the wells to hold the eggs in place. Embryo positions are adjusted with plastic forceps. 5) When the low-melt agarose sets, the live-imaging medium is added to the dish. *Right:* Schematic of embryo in Configuration 1. (B) *Configuration 2: Top loaded microwells for imaging with an inverted microscope.* 1) 700 μL of agarose is pipetted into the middle of the glass-bottom dish. The insert and slide are lowered onto it, taking care not to trap bubbles. 2) Agarose sets, and then the insert and slide are gently removed. 3) Embryos are loaded into microwells, as described above. 4) Live-imaging medium is added. *Right:* Schematic of embryo in Configuration 2.

**Figure 3.**
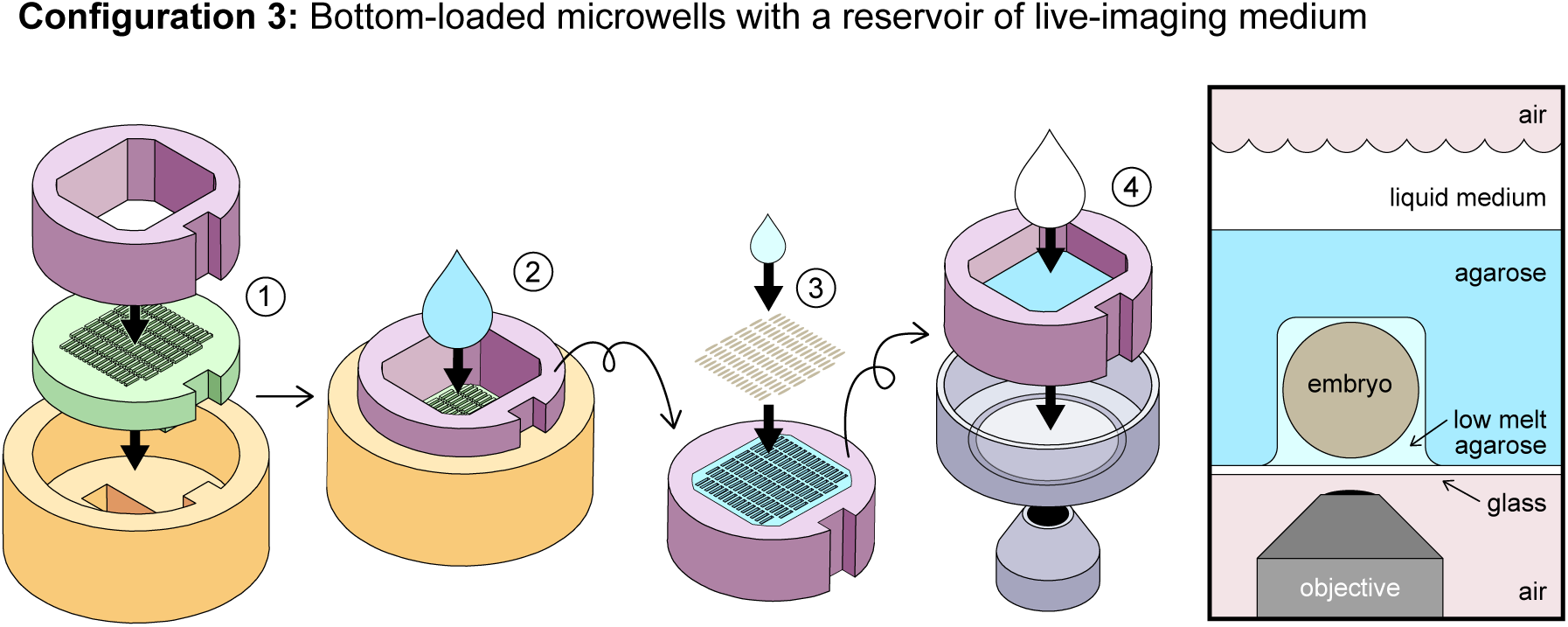
OMMAwell configurations for bottom-loaded microwells. *Configuration 3: Bottom loaded microwells with a reservoir of live-imaging medium.* 1) The insert (green) and cylinder (purple) are placed into the sheath (orange). 2) Molten agarose is poured into the cylinder to the desired depth. 3) Once the agarose has set, the cylinder and the agarose block are removed from the sheath and insert. The cylinder is flipped over, and the exposed microwells are loaded with embryos, as described in Figure 2. 4) The cylinder and agarose block are lowered into the glass-bottom dish, and live-imaging medium is added in the cylinder. *Right:* Schematic of embryo in Configuration 3.

### Configuration 1: Top loaded microwells for injecting or imaging with an upright microscope

(Figure 2A; see legend for step-by-step usage instructions). In this arrangement, the user can adjust the height of the mold insert, which is then lowered into molten 1.5% (w/v) agarose. Once the agarose has cooled and set, the mold is removed, leaving microwells in which to place the samples. Optionally, a small quantity of 0.7% (w/v) low-melt agarose (40-100 µL) can then be added to hold samples in the wells. When they are robjectives. It is also useful for holding samples that will be injected, such as with double-stranded RNA, small molecule activators or inhibitors, or recombinant protein (e.g. Donoughe et al., 2014). In this configuration, cricket embryos will successfully complete embryogenensis (~12 days), so long as the medium level is maintained. A drawback of this configuration is that the imaging medium may evaporate over the course of a time-lapse, and the lid cannot be added to prevent evaporation without blocking the microscope objective.

### Configuration 2: Top loaded microwells for imaging with an inverted microscope

(Figure 2B; see legend for step-by-step usage instructions). This is similar to Configuration 1, but now the mold insert is placed flat on molten 1.5% (w/v) agarose in a glass-bottom dish, producing microwells in a thin agarose film. The samples are loaded into the wells, fixed in place with 0.7% (w/v) low-melt agarose as above, the dish is covered with its lid, and then imaged from below on an inverted microscope. Because the lid remains on the dish and reduces evaporation, this configuration is the best one for long-term live imaging. As for configuration 1, but without the need to maintain the medium level manually, cricket embryos mounted in configuration 2 will develop healthily for 12 straight days, completing embryogenesis at rates comparable to unmounted embryos (92 to 100% hatching). A minor difficulty of this configuration may be that removing the mold insert and slide (Figure 2B, Step 2) without disrupting the thin agarose film is a delicate procedure for some insert designs. To ameliorate this problem, we reccomend use of 2% (w/v) agarose to make the microwells, and when it has set, pull up the mold insert with the agarose still adhered to it. Then, using plastic forceps, peel the agarose from the insert, return it to the glass-bottom dish, and “glue” it in place with ~100 µL of 0.7% low-melt agarose.

### Configuration 3: Bottom loaded microwells with a reservoir of live-imaging medium

(Figure 3; see legend for step-by-step usage instructions). This configuration is reccomended for cases where making the agarose film in Configuration 2 is troublesome for a particularly complex mold insert, or if it is necessary to use a larger volume of imaging medium than can be poured into the glass-bottom dish. The main advantage over Configuration 2 is that the mounting process is extremely robust. The downside is that the samples are separated from the imaging medium by a much thicker layer of agarose, which means that gas exchange is reduced. In our hands, cricket embryos mounted in this fashion will typically develop normally for 6 to 12 hours and then arrest. If the embryos are subsequently removed from their microwells and immersed in water, development continues. This configuration also offers a larger reservoir that can be filled and capped with a lid; its volume can be increased further by adding more layers to the “cylinder” in Step 5 of Supplemental Materials S3.

We have used each of these three configurations (Figures 2, 3) to live-image more than 100 embryos simultaneously. For species with smaller embryos, the maximum sample size is even larger. Since the array of wells is larger than the microscope’s field of view, a motorized stage is required to tile over the full set of embryos. (Alternatively, a user could manually move the stage in XY to image each embryo in the set. Because each well has a unique identifier, even with this manual approach, large numbers of individual embryos can be followed and uniquely identified over time-lapse periods.) As an example, we show a single time point from a time-lapse of 44 nuclear-marked transgenic cricket embryos (Figure 4A). We used a motorized stage to capture tiled micrographs of the full set of eggs once every five minutes. The recording continued for five days with no signs of phototoxicity or developmental defects. The specimens were then returned to the incubator, and 41 of the 44 embryos hatched.

**Figure 4.**
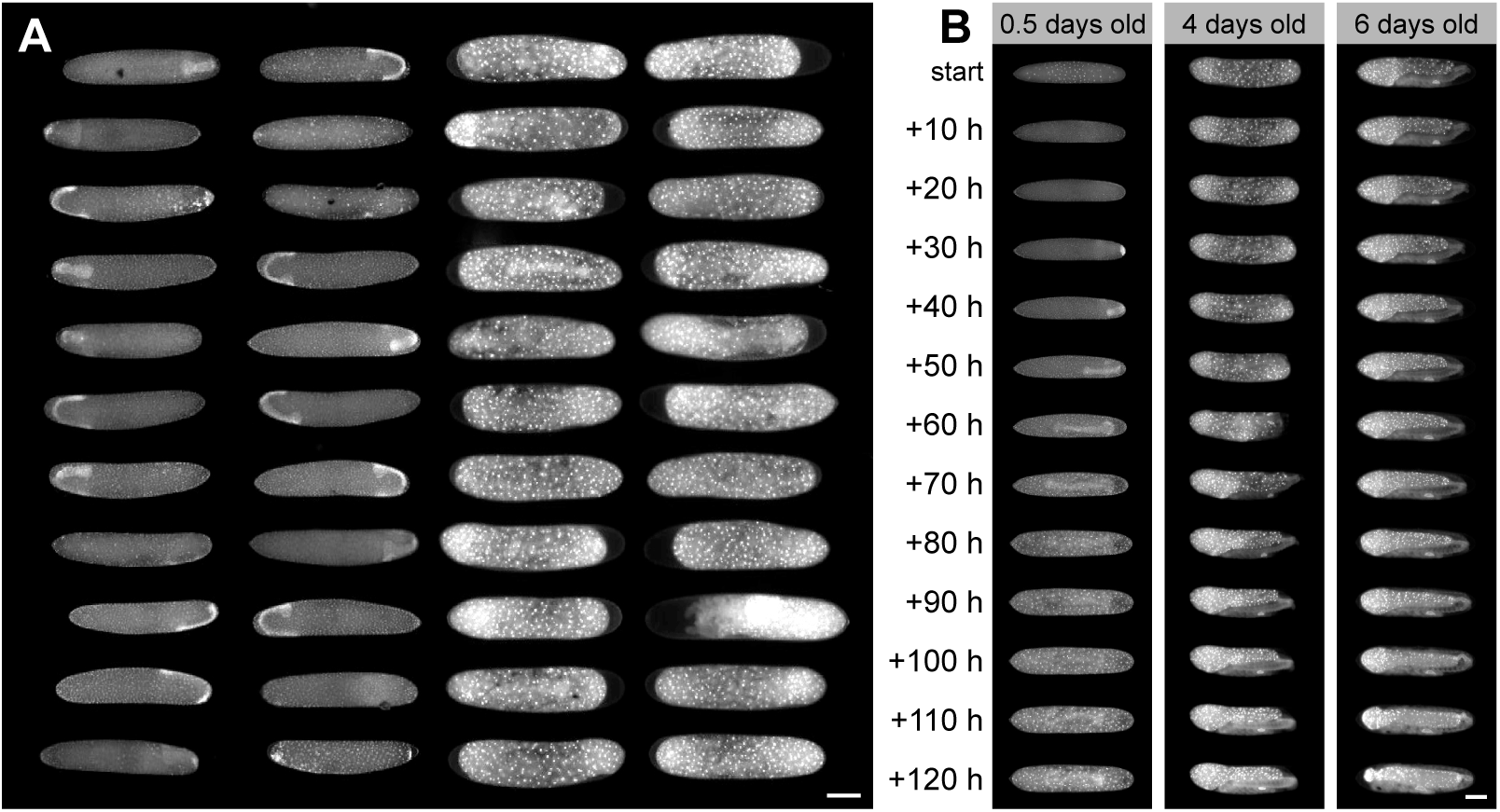
Arrays of live-imaged embryos within the OMMAwell mold. (A) A single timepoint from a time-lapse of an array of nuclear-marked transgenic cricket embryos in microwells. The two leftmost columns show germ band stage embryos that are beginning the physical re-orientation within the egg called anatrepsis. The two rightmost columns show a later stage when embryos are fully immersed in the yolk below the extraembryonic membrane called the serosa. (B) Time series of cricket embryos starting at different ages. The ambient temperature is ~24°C, so development is slower than that reported byDonoughe and Extavour (2016). Scale bars = 500 μm.

Researchers can easily design and fabricate their own mold inserts to generate wells of specified shapes, dimensions, and spacing. To make a new pattern, simply alter the design in one of the included insert DXF files (Supplemental Materials S2) and then engrave in acrylic using a laser cutter. Because these inserts are engraved in 2D, designs are limited to forms built from flat planes. However, if a more complex design were desired, such as a one with curved bottoms to the wells, the user could mill or 3D print it and mount it with the rest of the device.

## Materials and Methods

Design and assembly details are given in Supplemental Materials S1 and S2. When making the microwells, agarose (Bio-Rad #1613101) was dissolved at 1.5% weight/volume (w/v) in distilled water (or 2% for firmer molds). Then, eggs were held in microwells with low-melt agarose (Bio-Rad #1613112) at 0.7% (w/v) in distilled water. Tap water was used as a live-culturing solution for wild-type cricket eggs, but the user could also pour molds with agarose dissolved in a live-imaging buffer that is appropriate for their samples.

Wild-type eggs were imaged with transmitted white light on a Zeiss Lumar dissection microscope. For fluorescent imaging, recordings were taken of eggs from a transgenic line in which the cricket actin promoter drives expression of the cricket Histone-2B protein fused to Enhanced Green Fluorescent Protein (H2B-EGFP) (Nakamura et al., 2010). The 5x objective on a Zeiss Celldiscoverer 7 was used for imaging. The array of microwells was tiled with the motorized stage under the control of Zen software (Zeiss). A z-stack was captured at each position, then later combined using Zen’s “Extended Depth of Focus” (mode = “Contrast”).

## Acknowledgements

We are grateful to the Harvard Neuroengineering Core for fabrication assistance, to Sebastian Gliem (Zeiss) and Doug Richardson for help with high-throughput imaging at the Harvard Center for Biological Imaging. We also appreciated the feedback from our beta testers, several of whom provided photos of microwells in use: Austen Barnett, Leo Blondel, Eva Fast, Andrew Gehrke, Michael Brent Hawkins, Mara Laslo, Mark Martindale, Taro Nakamura, Megan Norris, Lorenzo Ricci, Elaine Seaver, Richard Smith, and Mansi Srivastava. For photo credits, please see the legends of Figure 1 and Supplemental Materials S1.

## Author Contributions

SD and CK designed and prototyped the tools with input from CGE. CK conducted brightfield microscopy. SD collected the time-lapse data. SD and CGE wrote the manuscript.

